# The unique nested gene *ALZAS* as a potential source of Aβ-peptides in Alzheimer’s disease

**DOI:** 10.1101/438630

**Authors:** Christin Gano, Heinz Reichmann, Bernd Janetzky

## Abstract

Amyloid-beta (Aβ) peptides are considered to be the cause of neuronal and synaptic cell death in Alzheimer’s disease (AD) since approximately 100 years. To date, it is assumed that Aβ-peptides arise from the proteolytic cleavage of the amyloid precursor protein (APP). However, within the *APP* gene, a nested gene called ALZheimer ASsociated (*ALZAS*) exists, which includes the entire Aβ_42_-sequence and thus may also be the origin of various Aβ-species. Here, we firstly confirmed expression and the postulated amino acid sequence of ALZAS and revealed the binding of selected monoclonal Aβ-antibodies to Aβ- and ALZAS protein. We confirmed the specificity of the anti-ALZAS antibody to ALZAS by protein sequence analysis. This anti-ALZAS antibody detects the same amino acid sequence as the autoantibody found in human blood of AD patients. Since no detailed data are currently available concerning *ALZAS* expression and the amount, localization and function of ALZAS protein in human cells and tissue, we performed gene (over)expression experiments on transcriptional and translational level. We verified a considerably lower mRNA amount of *ALZAS* compared to the host gene *APP*. Nevertheless, *ALZAS* transcription and translation seems to be heavily regulated in different human tissues and cells. Artificially increased mRNA levels of *ALZAS* did not led to an enhanced protein amount or considerable increase in cell death. Notably, cell localization of the ALZAS protein showed accordance to endosomes indicating that ALZAS, which contains the whole transmembrane domain of APP, might be a peripheral membrane protein. Since endosome dysfunctions are a characteristic event in early stages of AD and the highest ALZAS-autoantibody levels were already detected in early AD stages, *ALZAS* might play a crucial role in AD pathology and could possibly be a further diagnostic marker.

## Introduction

Alzheimer’s disease (AD) is the most common form of dementia marked by a progressive impairment of cognition, especially memory (Lambon Ralph et al. 2003). The significant loss of neurons (Hyman et al. 1984) and synapses (Scheff et al. 2006) is ascribed to pathological protein depositions called amyloid plaques (Terry et al. 1964). Amyloid plaques consist mainly of Aβ-peptides which are metabolites of the integral membrane protein APP (Selkoe et al. 1988). *APP* is ubiquitously expressed but it’s cellular function remains unclear, although studies suggest an involvement in cell adhesion (Sondag and Combs 2004; Sosa et al. 2013), memory and synapse formation (Mileusnic et al. 2000; Reinhard et al. 2005) and/or a regulative role in platelet function and immune response (Li et al. 1999). In the amyloidogenic pathway, APP is processed by β- and γ-secretases, leading to the formation of Aβ-peptides consisting of 40 or 42 amino acids (Chen and Schubert 2002; Edbauer et al. 2002). In AD patients, the amphiphilic Aβ-peptides are deposited as insoluble fibrils intraβ and/or extracellularly in the brain (LaFerla et al. 2007). According to the amyloid cascade hypothesis, these accumulated Aβ-peptides are neurotoxic (Hardy and Higgins 1992). However, the amyloid plaques are not necessarily associated with synaptotoxicity (Mucke et al. 2000) and can be found in the brain of non-demented patients too (Dickson et al. 1992; Nash 1997). In contrast, the Ae-derived diffusible ligands (ADDL) hypothesis reveals a stronger correlation between synaptic loss and soluble Aβ-peptides (Selkoe 2004). High concentrations of soluble Aβ-oligomers were detected in extracellular liquids, especially (C- and N-terminally) truncated Ar-peptides in cerebrospinal fluid of AD patients. The current doctrine states that all short isoforms of Aβ originate from APP (Portelius et al. 2011). However, there exists a unique nested gene called *ALZAS*, which contains the entire Aβ_42_-sequence as well (Figure 1).

*ALZAS* is localized within the *APP* gene on chromosome 21 (Kienzl et al. 2002) and represents a special form of overlapping genes called nested genes. These genes are embedded within the locus of a much bigger host gene (Gibson et al. 2005). Different from other nested genes, *ALZAS* possess the same reading frame as the host gene *APP* (Figure 1) (Kienzl et al. 2002).

Some nested genes coexisting on human chromosome 21 and 22 are associated with specific disorders (Karlin et al. 2002). Evidence for a correlation between *ALZAS* and AD lies in the existence of an endogenous autoantibody interacting with ALZAS (anti-ct12 IgG) identified in serum of AD patients (Bergmann et al. 2002). The highest antibody titers were measured in patients over 65 years suffering from depression or mild cognitive impairment. Non-demented controls exhibited no or lower concentrations of autoantibody. The 12 amino acids embodying the immunogenic sequence are located on the C-terminus of ALZAS (ct12). So far, no known isoform of APP or other proteins exhibit this sequence because it is encoded only by intron sequences of *APP* (Figure 1). Hence, our objective was to verify the specificity of the anti-ct12 ALZAS antibody and to examine co-detection of Aβ-antibodies to Aβ- and ALZAS protein. Further, we conducted *ALZAS* gene (over)expression experiments to: (I) compare transcript levels of *ALZAS* and *APP*, (II) reveal correlations between *ALZAS* expression and age, tissue/cell type or disease and (III) investigate the role of *ALZAS* in human cell metabolism by identification the localization of ALZAS in human cells.

**Figure 1.**
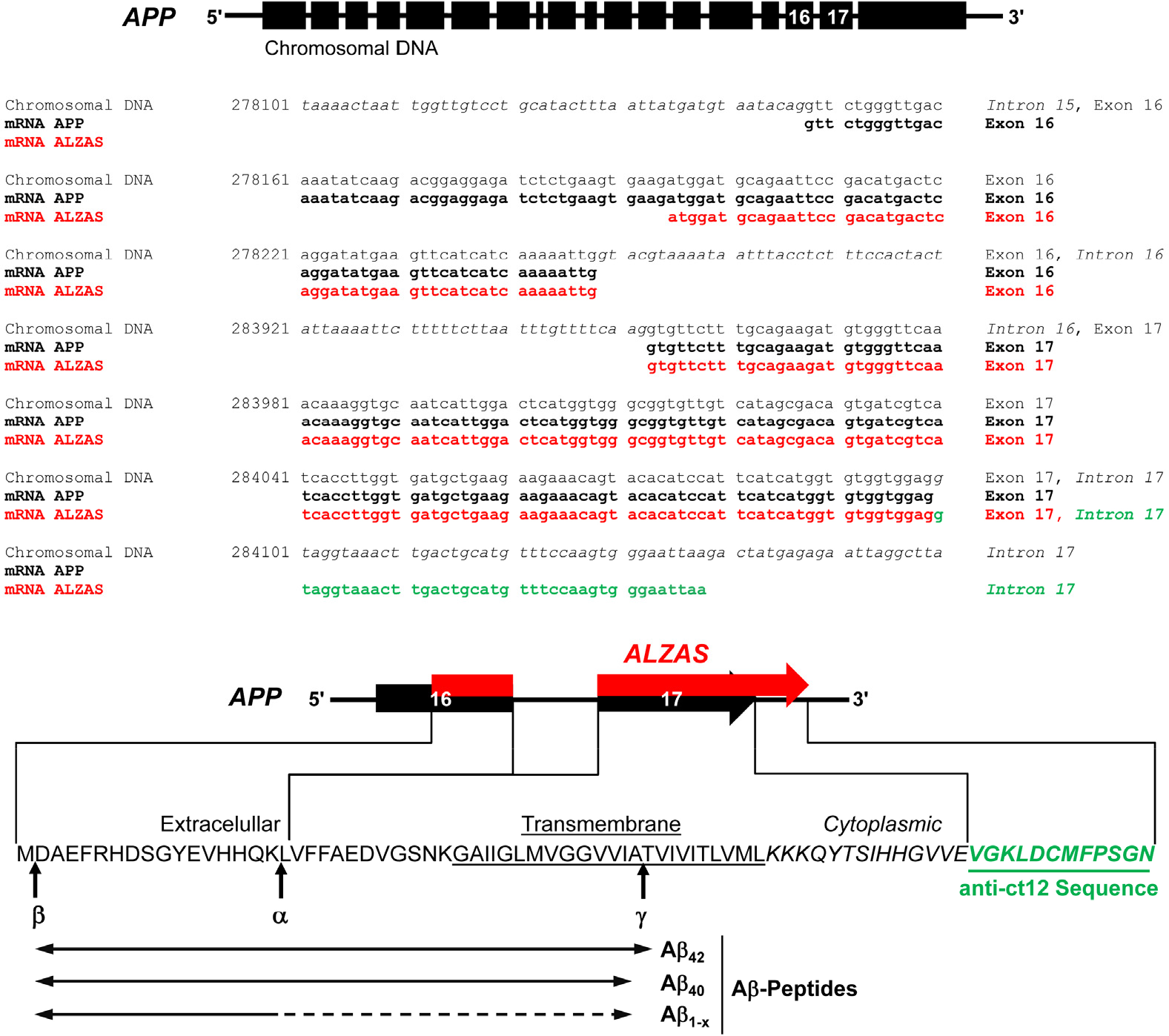
Molecular structure of the human *APP* gene and the nested gene *ALZAS*. The *ALZAS* gene is located within the human *APP* gene on chromosome 21. APP isoforms are encoded by up to 18 exons, while all include parts of the *ALZAS* gene. The coding sequence of *ALZAS* (red-green) starts in exon 16 of *APP* (NCBI accession number NG_007376, nucleotide 278195), excludes intron 16 and extends into intron 17 (NG_007376, nucleotide 284137). The derived amino acid sequence of ALZAS contains the whole Aβ_42_-sequence, cutting sites for α-, β- and γ-secretases and a C-terminal sequence of 12 amino acids (anti-ct12) derived from intron 17 (green; including stop codon) that is recognized by an endogenous autoantibody from AD patients. Postulated cell localization and Aβ-cleavage products of ALZAS are labeled.

## Materials and Methods

### Human tissue samples

All human tissue samples were from non-AD controls and AD patients differing in age and sex (Table S1). The tissue samples were obtained post mortem (post mortem time <24 hrs, kindly provided by Prof. Jellinger, University of Technology, Vienna), postnatal, in the lifetime or from transplant following informed consent of the subjects. All samples were stored at −80°C until required. Working with human tissue samples were in accordance with the Helsinki convention and approved by Ethical Committee of the University of Dresden (EK197092004).

### Total RNA isolation

Total RNA was isolated from human tissue using RNeasy Lipid Tissue Mini Kit (Qiagen) or PAXgene Blood RNA Kit (PreAnalytix). Total RNA from human neuroblastoma SH-SY5Y cells was isolated with RNeasy Mini Kit (Qiagen) in accordance with the manufacturer’s instructions. Details regarding RNA integrity are provided in the Supplementary section “RNA analysis”.

### Protein extraction

#### Human tissue (Total protein isolation)

Proteins from human tissue (Table S1) were extracted using a tissue grind and protein lysis buffer comprised of PBS, 0.5% v/v Triton X-100 (both Thermo Fisher Scientific), 0.2% v/v SDS (Carl Roth) and protease inhibitors (Sigma-Aldrich or Roche). The tissue lysate was sonicated for 5 min and afterwards incubated for 5 min at 99°C. The tissue lysate was centrifuged for 5 min at 4°C and 20,000 *g*. The supernatant was collected and stored at ≤−20°C until required.

#### Fractionated protein isolation (Membrane protein protocol)

The tissue sample of a non-demented patient (Cortex, sample 118; Table S1) was homogenized in 300 μl protein lysis buffer and centrifuged for 5 min at 4°C and 900 *g*. Supernatant (s1, water-soluble and membrane proteins) was collected and again centrifuged for 30 min at 4°C and 230,000 *g*. The supernatant (s2, water-soluble proteins) was separated and the pellet (membrane proteins) was resuspended in 200 μl protein lysis buffer. All samples were stored at ≤−20°C until usage.

#### Adherent cells

To isolate proteins from adherent human neuroblastoma cells SH-SY5Y, the medium was removed and the cells washed with PBS. To lyse the cells, protein lysis buffer was added. A modified protocol was applied where secreted protein, protein being part of death (detached) cells and protein in attached cells (cell extract) required separation. For that, the medium was centrifuged for 5 min at 14,000 *g* and the supernatant (secreted proteins) and the death cells were separated. The death cells were resuspended in protein lysis buffer. All lysates were stored at ≤−20°C.

### Gene expression (mRNA) analysis using RT-qPCR

1-step qPCR was performed with Brilliant II SYBR Green QRT-PCR Master Mix Kit (Agilent Technologies) according to manufacturer’s instructions. For 2-step qPCR, total RNA was first reverse transcribed with QuantiTect Reverse Transcription Kit (Qiagen) to generate cDNA and afterwards amplified using QuantiTect SYBR Green PCR Kit (Qiagen) following the manufacturer’s instructions. All qPCRs were performed in a Mx3000P cycler (Agilent Technologies). Cycling conditions for 1-step RT-qPCR: 25 min 50°C / 15s 95°C / 45 cycles of 15s 94°C, 20s 57°C, 30s 72°C (modified cycling conditions comprised an additional temperature plateau for 15s at 77°C) / 1 cycle of 1 min 95°C,
 30s 55°C, 30s 95°C. Cycling conditions for 2-step RT-qPCR: 15 min 95°C / 40-45 cycles of 15s 94°C, 30s 57°C, 30s 72 °C / 1 cycle of 1 min 95°C, 30s 55°C, 30s 95°C. In 1-step RT-qPCRs, each primer was applied at a final concentration of 0.1 pmol/μl and in 2-step RT-qPCRs at 0.15 pmol/μl. Each primer pair was tested for their efficiency using a RNA or cDNA serial dilution (1-step RT-qPCR: 0.2-200 ng RNA; 2-step RT-qPCR: 0.001-2 μl cDNA).

Each RNA sample was performed in duplicate to quadruplicate in qPCR and analyzed using MxPro software (Agilent Technologies). Samples were normalized to *GAPDH* (glyceraldehyde-3-phosphate-dehydrogenase) and *HMBS* (hydroxymethylbilane synthase) (Vandesompele et al. 2002; Gutala and Reddy 2004) according to the relative quantification method (Bustin 2004). SH-SY5Y cell samples were further normalized to controls, deducting the non-induced expression level.

Relative changes in gene expression levels (x-fold) were calculated according to −2^ΔΔc_T_^ (Livak and Schmittgen 2001). Dixons Q-Test was applied to exclude extraneous values to 99% significance (Dean and Dixon 1951). Tissue RNAs that were heavily degraded were excluded from RT-qPCR experiments.

### (Over)expression of the *ALZAS* gene in the human neuroblastoma cell line SH-SY5Y using the Regulated Mammalian Expression System (Promega)

The human neuroblastoma cells SH-SY5Y (LGC Standards) were first stably transfected with the regulator vector pReg neo (Promega) (SH-SY5Y-pReg neo) and, subsequently, transient transfected with the expression plasmid pF12A-ALZAS (SH-SY5Y-pReg neo-pF12A-ALZAS). Stable transfection of pReg neo was performed after vector linearization. For that, pReg neo was treated for 3 hrs with the restriction enzyme *Xmn*I according to the instruction manual (Thermo Fisher Scientific) and afterwards purified using the QIAquick Gel Extraction Kit (Qiagen).

The expression plasmid pF12A-ALZAS (clone 2, Supplementary Figure S1a) was generated by cloning of human *ALZAS* gene sequence into the expression vector pF12A RM Flexi (Promega). For detailed information regarding cloning process, see Supplementary section “Plasmids”. Cell culture details are depicted in Supplementary section “Adherent Cell culture of human neuroblastoma cells SH-SY5Y”.

48 hrs after transfection, SH-SY5Y-pReg neo cells were selected with geneticin disulfate (G418) solution (Carl Roth). For re-cloning, the cells were treated with 10% FBS/DMEM medium comprised of 88.8% v/v Dulbecco’s modified eagle medium (DMEM, 1 g/l D-glucose, Thermo Fisher Scientific) supplemented with 10% v/v fetal bovine serum (FBS, Sigma-Aldrich), 1% v/v hepes buffer (Sigma-Aldrich) and 0.2% v/v penicillin-streptomycin (500x, Roche). After 7 days of cultivation, G418 was added again. To confirm the integration of pReg neo in the genome of SH-SY5Y, total RNA was isolated as described above followed by 2-step RT-qPCR with the primer pair pReg neo-f1 and pReg neo-r1 (Table S2).

The appropriate G418 concentration for selection of SH-SY5Y-pReg neo cells (Southern and Berg 1982) was determined. For that, the SH-SY5Y cells were treated with varying concentrations of G418 from 0 μg/ml (control) up to 1000 μg/ml over a time interval of 1-12 day(s). The viability of the adherent cells was investigated by a live/dead staining with propidium iodide (Supplementary Figure S2b). Finally, SH-SY5Y-pReg neo cells were maintained at 600 μg/ml G418 concentration. The second transfection of the pF12A plasmid pF12A-ALZAS, pF12A-ALZASinv or pF12A-hRluc (the latter two serves as control) was performed as described below and before every new gene expression experiment. The control plasmid pF12A-ALZASinv was utilized in case of ALZAS protein analysis, pF12A-hRluc (kindly provided from Promega) for the analysis of different SH-SY5Y-pReg neo clones and water (no-template) was transfected as a control for *Renilla* Luciferase Assay or *ALZAS* mRNA verification. For detailed information regarding measurement of luciferase activity see Supplementary section “Luciferase assay”.

To induce (over)expression of the gene cloned into the pF12A plasmid, the SH-SY5Y-pReg neo cells were treated with coumermycin (Promega) after transfection mixture was removed. The inductor was diluted to the desired concentration (7-500 nM) in 10% FBS/DMEM medium containing G418. Controls transfected with pF12A-ALZASinv or sterile water (no-template) were also treated with coumermycin. To determine the basal expression level of *ALZAS* or *Renilla* luciferase, one well of cells was not treated with coumermycin (= non-induced SH-SY5Y-pReg neo-pF12A cells). Supplementary Figure S2a illustrates a schematic timetable for the *ALZAS* gene (over)expression studies in the human neuroblastoma cell line SH-SY5Y.

### Plasmid transfection into SH-SY5Y

Prior to transfection procedure, the SH-SY5Y cells were plated at a density of 100,000 cells/cm^2^. One day later, the plasmid was inserted into SH-SY5Y under sterile conditions using the Lipofectamine 2000 Kit according to manufacturer’s instructions (Thermo Fisher Scientific). The cells were incubated with the transfection mixture for 3 hrs (for pF12A-ALZAS, pF12A-hRluc, pEGFP-ALZAS, pEGFP-C1) or 6 hrs (only for linearized pReg neo or pF12A-ALZAS) at 37°C, 5% CO_2_ and saturated humidity. The mixture was removed and replaced by 10% FBS/DMEM medium.

### In vitro translation

For *in vitro* translation, a DNA construct was synthesized comprised of a T7 promoter followed by the coding sequence *EGFP* and *ALZAS* and a poly(A) signal. For that, a PCR using the primers T7pA-GFPA-f and T7polyA-A-r (Table S2) was performed, with the plasmid pEGFP-ALZAS clone 3 (Supplementary Figure S1b) as template DNA. Details regarding cloning of human *ALZAS* gene into the vector pEGFP-C1 are provided in Supplementary section “Plasmids”. The Platinum *Pfx* DNA Polymerase PCR mixture was modified for 10x *Pfx* Amplification Buffer (2-fold), dNTP (0.2 mM each), primer (0.2 pmol/μl each) and cycling conditions: 5 min 94°C / 10 cycles of 15s 94°C, 20s 50°C, 10s 68°C / 25 cycles of 15s 94°C, 20s 57°C, 10s 68°C / 5 min 68°C. A control DNA construct with *Firefly* luciferase (Fluc) was also amplified employing the primers T7polyA-Fluc-f, T7polyA-Fluc-r (Table S2) and as template a luciferase control DNA (Promega). Cycling conditions were as follows: 5 min 94°C / 35 cycles of 15s 94°C, 20s 59°C, 25s 68°C / 5 min 68°C. Both PCR products (T7-EGFP-ALZAS-polyA, T7-EGFP-Fluc-polyA) were purified using QIAquick Gel Extraction Kit and the DNA sequence analyzed (T7-EGFP-ALZAS-polyA: Supplementary Figure S1c, T7-EGFP-Fluc-polyA: DNA sequencing data not shown).

To synthesize proteins (EGFP-ALZAS or Fluc) from the PCR-generated DNA templates, either the TNT T7 Quick for PCR DNA System or the TNT T7 Quick Coupled Transcription/Translation System (both incorporated in the T7 Sample System, Promega) with Transcend tRNA (Promega) was employed following instruction manuals. For detailed information regarding measurement of luciferase activity see Supplementary section “Luciferase assay”.

### Immunoprecipitation

The *in vitro* translation product EGFP-ALZAS was mixed with GFP-binding agarose beads (Chromotek gta) according to the instruction manual and incubated for 1 hr at 4°C. Dilution buffer was supplemented with 1 mM phenylmethylsulfonyl fluoride (Roth) and protease inhibitor. The beads were washed three times with ice cold PBS and finally resuspended in 20 μl 1-fold laemmli buffer (62.5 mM Tris-HCl pH 6.8, 12% w/v SDS, 60% v/v glycerin (all Carl Roth), bromophenol blue (Sigma-Aldrich), freshly added 0.05 M DTT (Carl Roth)). After heating and centrifugation, the supernatant was removed (bound fraction with EGFP-ALZAS fusion protein). The beads were treated once more with 20 μl 1-fold laemmli buffer. The bound fractions were merged and either stored at −20%C until usage or the whole volume applied in a SDS-PAGE immediately. SDS gel was stained with Coomassie dye prior to protein sequence analysis. Staining procedure was carried out for 1 hr in 0.2% w/v Coomassie Brilliant Blue R (Sigma-Aldrich), 10% v/v acetic acid (Merck) and 50% v/v methanol (VWR). The gel was destained four times for 15 min in 2% acetic acid and 49% methanol. After destaining overnight, the gel was rinsed with water four times for 15 min and stored a 4%C prior to protein sequencing at the Max Planck Institute of Molecular Cell Biology and Genetics in Dresden (Mass Spectrometry facility).

### Mass spectrometry-based protein sequencing of EGFP-ALZAS fusion protein

Coomassie stained band was excised and its protein content was digested overnight with trypsin. The resulting peptide mixture was extracted twice with 5% formic acid and acetonitrile and analyzed by liquid chromatography tandem mass spectrometry (LC MS/MS) on an Ultimate3000 nanoLC system interfaced on-line to a LTQ Orbitrap Velos hybrid tandem mass spectrometer (both Thermo Fisher Scientific) as described (Vasilj et al. 2012). Peptides were matched by MASCOT v.2.2.04 software (Matrix Sciences) search against the EGFP-ALZAS construct sequence.

### SDS-PAGE

Proteins were separated in a SDS-PAGE (Laemmli 1970). The protein concentration in tissues or cell lysates was measured using the Quant-iT Protein Assay Kit and the Qubit Fluorometer according to the instruction manual (Thermo Fisher Scientific). 30 μg of total protein amount, 10 μl of immunoprecipitation (IP) fractions or 2-4 μl *in vitro* translation product was prepared in laemmli buffer (final buffer concentration 1-fold). Protein samples were incubated for 10 min at 95%C followed by a centrifugation step for 2 min at 10,000 *g* (2,500 *g* for bound fraction of IP). The protein samples were then loaded on a 14% Tris-glycine gel (Anamed) and separated by electrophoresis for 1 hr 45 min at 25 mA per gel and 125 V in Tris-glycine-SDS buffer (1.92 M glycine, 0.25 M Tris, 1% w/v SDS (all Carl Roth); buffer was filtered when using IP protein probes). A prestained protein marker (Thermo Fisher Scientific) was used to identify protein sizes. For tissue and cell probes, the antigen ALZAS ct12 peptide (<10 kDa, Davids Biotechnology, Regensburg, Germany) was loaded as a control on the gel.

### Immunoblotting

After SDS-PAGE, the protein samples were transferred to nitrocellulose membrane using iBlot Dry Blotting System according to the instruction manual (blotting time 8 min) (Thermo Fisher Scientific). The membrane was blocked for 1-2 hrs at room temperature in 5% w/v skim milk powder (Carl Roth) in PBST (PBS, 0.1% v/v Tween 20 (Serva Electrophoresis)). PBS was used for washing: 1.37 M NaCl, 81 mM Na_2_HPO_4_, 14.7 mM KH_2_PO_4_ (all Merck), 27 mM KCl (Sigma-Aldrich) and filtered when IP protein probes were analyzed.

After blocking, the membrane was incubated overnight at 4%C with primary antibody diluted in 1% w/v skim milk powder in PBST. Primary antibodies used are: anti-ALZAS (1:500-1:1000, polyclonal, rabbit, affinity purified IgG, Davids Biotechnology, Regensburg, Germany), anti-GAPDH (1:700,000, Merck MAB374), anti-GFP (1:2000, Abcam ab290), anti-beta Amyloid 1-13 (1:500, monoclonal, Abcam ab17331), anti-beta Amyloid 22-35 (1:5000, polyclonal, Abcam ab62658) or anti-beta Amyloid 1-14 (1:1000, polyclonal, Abcam ab2539). The membrane was washed three times for 10 min with PBST. Then, the membrane was incubated 1-2 hrs at room temperature with a secondary antibody diluted in 1% w/v skim milk powder in PBST. Secondary antibodies were conjugated to HRP (1:5000, Jackson ImmunoResearch anti-mouse 715-035-150 or anti-rabbit 711-035-152). The membrane was washed again three times for 10 min with PBST. Protein samples on the membrane were visualized with ECL Western Blotting Detection Reagents and Analysis System (GE healthcare) by measuring the chemiluminescence in the Luminescent Image Analyzer LAS-300 (Fujifilm; 5 min acquisition time). Each tissue protein sample was performed at least in duplicates (separate blots) and their protein signals were quantified using ImageJ software.

### Immunocytochemistry and fluorescence microscopy

For immunocytochemical staining, we used following adherent cells: SH-SY5Y, human fibroblasts, dopaminergic neurons (cell line T12.9, differentiated 24 days following Reinhardt et al. 2013) (the last two are kindly provided by Prof. Storch, Department of Neurology, University of Rostock). The cells were rinsed with PBS and fixed for 10 min at room temperature with 4% w/v paraformaldehyde (Sigma-Aldrich), 96% PBS (pH 7.4). The cells were rinsed three times with PBS. For PDI or F-Actin staining, the cells were incubated for 1-2 hrs at room temperature with freshly prepared blocking solution consisting of PBS-Triton X-100 (0.2% v/v Triton X-100 in PBS) and 3% v/v donkey serum (Jackson ImmunoResearch). Otherwise, cells were first permeabilized for 10 min at room temperature in PBS-Triton X-100 and blocked for 1-2 hrs at room temperature in 93% PBS, 5% v/v donkey serum and 2% w/v bovine serum albumin (BSA, Serva Electrophoresis albumin bovine fraction V). Primary antibody incubation was carried out overnight at 4%C. Used antibodies were diluted in blocking solution: anti-ALZAS (1:100, polyclonal, rabbit, affinity purified IgG, antigen ct12, Davids Biotechnology, Regensburg, Germany), anti-APP (1:100, Thermo Fisher Scientific 13-0200), anti-PDI (1:100, Enzo Life Sciences ADI-SPA-891), anti-GM130 (1:400; BD Biosciences 610822), anti-Cytochrome C (1:400, BD Biosciences 556432), anti-Cathepsin (1:400, Santa Cruz sc-6499), anti-Rab5 (1:200, Abcam ab50523), anti-Rab7 (1:200, Abcam ab50533) or anti-GFP (1:500, Abcam ab13970). After washing four times for 10 min with PBS, cells were incubated for 1 hr in the dark with secondary antibodies conjugated with a fluorescent dye. The following antibodies were diluted in blocking solution: anti-rabbit IgG Alexa Flour^®^ 488 (1:500, A-21206), anti-rabbit IgG Alexa Flour^®^ 555 (1:500, A-31572), anti-mouse IgG Alexa Flour^®^ 488 (1:500, R37114), anti-mouse IgG Alexa Flour^®^ 555 (1:500, A-31570), anti-goat IgG Alexa Fluor^®^ 488 (1:500, A-11055), Rhodamine phalloidin (1:100, R415) (all Thermo Fisher Scientific) or anti-chicken IgY, FITC conjugate (1:500, Merck Millipore AP194F). The cells were rinsed four times for 10 min with PBS. Nuclear DNA was mounted in the dark for 4 min in PBS, 0.075% v/v Hoechst 33342 (10 mg/ml, Thermo Fisher Scientific). The cells were again rinsed four times with PBS. Stained cells attached on coverslips were mounted with Fluoromount-G (Southern Biotech) on slides.

Stained cells were imaged using fluorescence microscope DMIRE2 (Leica) and the Leica software FW 4000 v1.1SP1. A 20x objective was applied and the emitted light was collected with 405, 488 and 594 nm filters. The images were analyzed in ImageJ software. Confocal microscopy was executed at MTZ Imaging Light Microscopy Facility (TU Dresden). The microscope TCS SP5 (Leica) and the LAS AF software (Leica) was utilized to image cell staining. A HCX PL APO lambda blue 63x 1.4 oil objective and the filter wavelengths of 415-475 nm, 498-533 nm and 553-700 nm were applied. Confocal images were further evaluated with the arivis browser x64 software.

### Cell viability tests

#### Live cell imaging of adherent cells

Adherent SH-SY5Y cells were maintained in a small incubator (37%C, 5% CO_2_, saturated humidity) connected to the fluorescence microscope DMIRE2 (Leica). Using Leica FW 4000 v1.1SP1 software and time-lapse microscopy, cells were recorded in phase contrast images every 15 min over a time period of 24 hrs.

#### Live/dead staining of adherent cells with propidium iodide

The medium was removed under sterile conditions and the adherent SH-SY5Y cells rinsed with PBS. 500 μl/2 cm^2^ of freshly prepared staining solution was added to the cells and incubated for at least 6 min at 37%C. Staining solution consisted of 20% v/v DMEM (1 g/l D-glucose), 0.08% v/v propidium iodide (100 μg/ml) and 0.04% v/v Hoechst 33342 (10 mg/ml) in PBS (all from Thermo Fisher Scientific). Cells were visualized on the fluorescence microscope DMIRE2 with the Leica FW 4000 v1.1SP1 software. Emission light filter wavelengths of 405 nm (Hoechst) and 594 nm (propidium iodide) were applied. Counted dead SH-SY5Y-pReg neo-pF12A-ALZAS cells (induced) were normalized to controls (mean of dead cells from no-template control, SH-SY5Y, SH-SY5Y-pReg neo-pF12A-ALZASinv, SH-SY5Y-pReg neo-pF12A-hRluc - each treated or not treated with coumermycin) and the value of non-induced SH-SY5Y-pReg neo-pF12A-ALZAS cells was deducted.

### Statistical analysis

Data are expressed as mean ± standard deviation. All statistical analyses were performed and figures created using Excel or GraphPad Prism 5.0 software.

## Results

### *ALZAS* gene expression studies in human tissues

In initial expression experiments, mRNA of the nested gene *ALZAS* was verified in frontal cortex of non-demented controls and AD patients by reverse transcriptase-quantitative real-time PCR (RT-qPCR) (Kienzl et al. 2006). So far, no data exist as to whether *ALZAS* transcription and translation show tissue- and/or disease-specific variations. Therefore, we first investigated organs differing in structure and function from persons of divergent age and sex. qPCR amplification of *ALZAS* mRNA was performed using primer pairs with reverse primers being located within an intron of *APP* (Figure 2a). On the other hand, the *APP* primer was designed to not detect *ALZAS*. All applied primer pairs exhibited PCR-efficiencies between 95-105%.

The analysis of different brain tissues (cortex (gray and white matter), occipital lobe, brainstem, hippocampus, precentral gyrus, postcentral gyrus) and the peripheral organs (heart, liver, spleen, skeletal muscle, blood and placenta) revealed an overall weak expression of *ALZAS* independent of sex (selected tissues shown in Figure 2b-d). A considerable difference in the mRNA level of *ALZAS* in the brain of non-AD controls and AD patients was not observed (Figure 2b), confirming previous results from our group (Kienzl et al. 2007). Nevertheless, a tissue-specificity in the *ALZAS* expression could be noticed (Figure 2b, c). While our investigations suggest a high *ALZAS* mRNA level in the heart of an AD patient (Figure 2c), the lowest *ALZAS* mRNA amount was verified in blood of children (Figure 2b). Comparing the expression of the nested gene *ALZAS* with its host gene *APP*, a consistent difference was revealed (Figure 2d). In almost all the tissues tested, the expression of *ALZAS* was nearly 1000-fold lower than the expression of *APP*.

In Western blot analysis, we detected protein bands interacting with anti-ALZAS antibody at approximately 60 kDa in the brain (Figure 2e), but in the peripheral tissues, heart and liver, we could observe additional bands at about 10 and 30 kDa (Figure 2f). In the placenta samples, only protein bands at about 10 and 30 kDa were identified (data not shown). Depending on the tissue type the protein bands exhibit distinct intensities. While no signals could be detected in spleen and skeletal muscle, the strongest signals were observed in the liver of an AD patient (Figure 2f). Divergent signal intensities were found in brain (Figure 2e). Obviously weaker signals were noticed in cortex in comparison to the hippocampus. Further, no evident differences could be observed between non-AD controls and AD patients in cortex and hippocampus.

**Figure 2.**
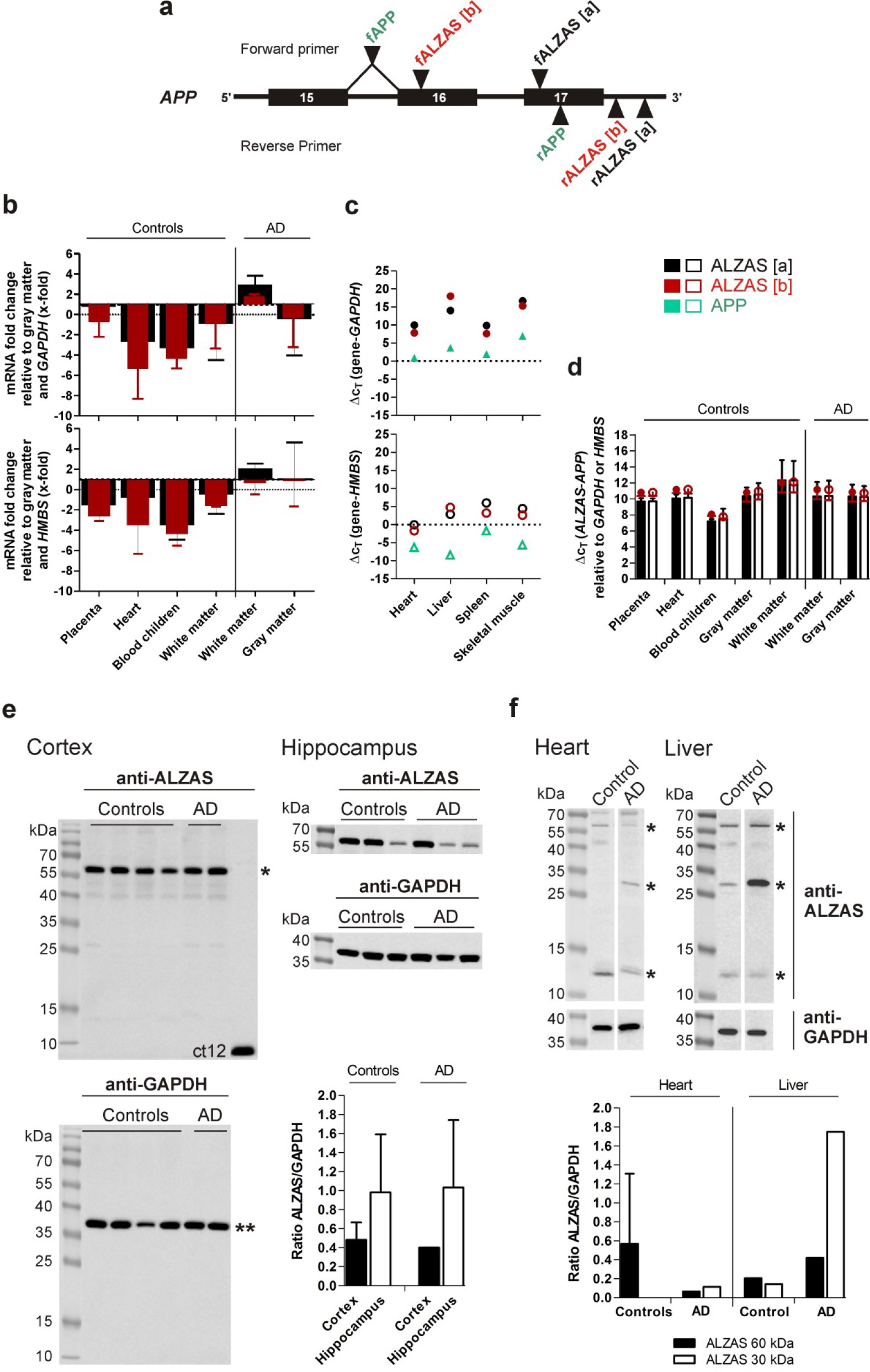
Measurement of *ALZAS* mRNA and protein in different human tissues. (a) Partial *APP* gene structure (exon 15 until intron 17) and location of primer pairs used for RT-qPCR amplification of *ALZAS* and *APP* mRNA. *ALZAS* mRNA was detected with two primer pairs of different amplicon size [a] = 173 bp and [b] = 236 bp. The amplicon of *APP* exhibited a size of 174 bp. f: Forward primer, r: Reverse primer. (b) Relative changes of *ALZAS* expression compared to cortex gray matter of non-AD controls (n = 11) (2-step RT-qPCR, normalized to *GAPDH* and *HMBS*), where a higher *ALZAS* mRNA level is illustrated by bars above 1 and a lower mRNA level by values below 1. Black bars illustrate *ALZAS* expression measured with primer pair [a] and red bars with primer pair [b]. Data is shown as mean ± SD determined from at least duplicate measurements. Placenta (non-AD) n = 10, gray matter (AD) n = 7, heart (non-AD) n = 6, blood children (non-AD), white matter (non-AD) and white matter (AD) n = 4. (c) Relative level of *ALZAS* mRNA in peripheral tissues of one AD patient (1-step RT-qPCR). Filled symbols are normalized to *GAPDH*, open symbols to *HMBS*. Black and red symbols illustrate *ALZAS* expression measured with primer pairs [a] or [b], respectively, green symbols represent *APP* expression. n = 1 per tissue. (d) Relation of *ALZAS* and *APP* gene expression in different tissues analyzed by 2-step RT-qPCR (normalized to *GAPDH* (filled symbols) and *HMBS* (open symbols)). Bars illustrate *ALZAS* expression measured with primer pair [a], red circles represent *ALZAS* mRNA measurement using primer pair [b]. Data is presented as mean ± SD determined from at least duplicate measurements. Placenta and gray matter (both non-AD) n = 11, heart (non-AD) and gray matter (AD) n = 7, white matter (AD) n = 5, blood children and white matter (both non-AD) n = 4. (e, f) Tissue lysates from brain and peripheral tissues were immunoblotted for ALZAS and GAPDH. Representative blots are shown. For better illustration, vertical blot lanes were partially rearranged. Lanes originated by different blots are separated. Asterisks indicate protein bands interacting with anti-ALZAS antibody (labeled as *) at about 10, 30 and 60 kDa or with anti-GAPDH (labeled as **) at 36 kDa. The ratio of the chemiluminescence signals from ALZAS and GAPDH were calculated. Data is shown as mean ± SD determined from at least triplicate measurements. Heart (non-AD) n = 8, cortex (non-AD) n = 4, hippocampus (non-AD) and hippocampus (AD) n = 3, cortex (AD) n = 2, liver (non-AD), heart and liver (both AD) n = 1. *GAPDH*: Glyceraldehyde-3-phosphate-dehydrogenase, *HMBS*: Hydroxymethylbilane synthase.

### Analysis of *ALZAS* gene (over)expression in human neuroblastoma cells SH-SY5Y

To date, nothing is known about the role and importance of *ALZAS* in human cell metabolism. Thus, we performed cell culture experiments in the human neuroblastoma cell line SH-SY5Y to examine the effects of regulated *ALZAS* (over)expression. To this end, we cloned the *ALZAS* gene in the controllable expression vector pF12A RM Flexi (Supplementary Figure 1a) which is part of a regulated expression system (Zhao et al. 2003). The second part of this system consists of the regulator vector pReg neo which allows for concentration-dependent gene expression of proteins potentially toxic to cells.

We first introduced the regulator vector pReg neo into the genome of SH-SY5Y cells to control the influence of the site of genome integration on basal (non-induced condition) and increased (induced) expression of *ALZAS*. 12 different neomycin resistant clones (SH-SY5Y-pReg neo) were transfected with the reporter plasmid pF12A-hRluc and several conditions, like different inductor concentrations and induction periods, were tested to identify clones with a low basal expression and a high expression under inducing conditions (Figure 3a). The two clones, no. 2 and 8, showed a low basal *Renilla* luciferase activity (Figure 3a), but differed by a factor up to 20 when comparing to the induced expression (Figure 3b). Both clones were chosen for subsequent *ALZAS* (over)expression studies on transcriptional (mRNA analysis by qPCR) and translational (protein analysis by immunoblotting, analyzing cell viability with live/dead staining) level.

To analyze *ALZAS* (over)expression, we transfected the SH-SY5Y-pReg neo clones 2 and 8 with the pF12A-ALZAS plasmid. Herein, we verified that a 12 hrs induction period yielded the highest *ALZAS* mRNA levels for both clones (Figure 3c). For clone 2, concentration of 200 nM was optimal to achieve highest *ALZAS* mRNA levels (>200-fold increase compared to controls). For clone 8, 10 nM coumermycin was ideal to obtain a >500-fold increase in *ALZAS* gene expression. Induction intervals above 24 hrs (36, 48 hrs) or an extended transfection duration of 6 hrs led to no further increase of *ALZAS* mRNA level in cells of SH-SY5Y-pReg neo-pF12A-ALZAS clone 2 (data not shown).

Western blot analyses with extracts from *ALZAS* overexpressing cells revealed protein bands interacting with anti-ALZAS antibody in the cell extract and in the supernatant (Figure 3e, f) but not in the dead cell fraction (detached cells, data not shown). In the cells of SH-SY5Y-pReg neo-pF12A-ALZAS clone 2 and 8, protein bands interacting with anti-ALZAS antibody were detected at about 30, 38, 60 and/or 100 kDa band, while in the supernatant, only a 38 kDa protein band was clearly visible. However, no increased signals were observed either for clone 2 or clone 8 (Figure 3e, f lane VII) when compared to non-induced SH-SY5Y-pReg neo-pF12A-ALZAS cells or controls. For clone 2, other tested conditions like timespans (24 or 48 hrs) or inductor concentrations (Supplementary Figure 2c lane VIII, IX) showed the same result. Moreover, after an expression period of 24 hrs (3 hrs transfection duration), no considerable increase in cell death was verified in SH-SY5Y-pReg neo-pF12A-ALZAS clone 2 and 8 via live cell imaging (data not shown) and live/dead staining (Figure 3d).

**Figure 3.**
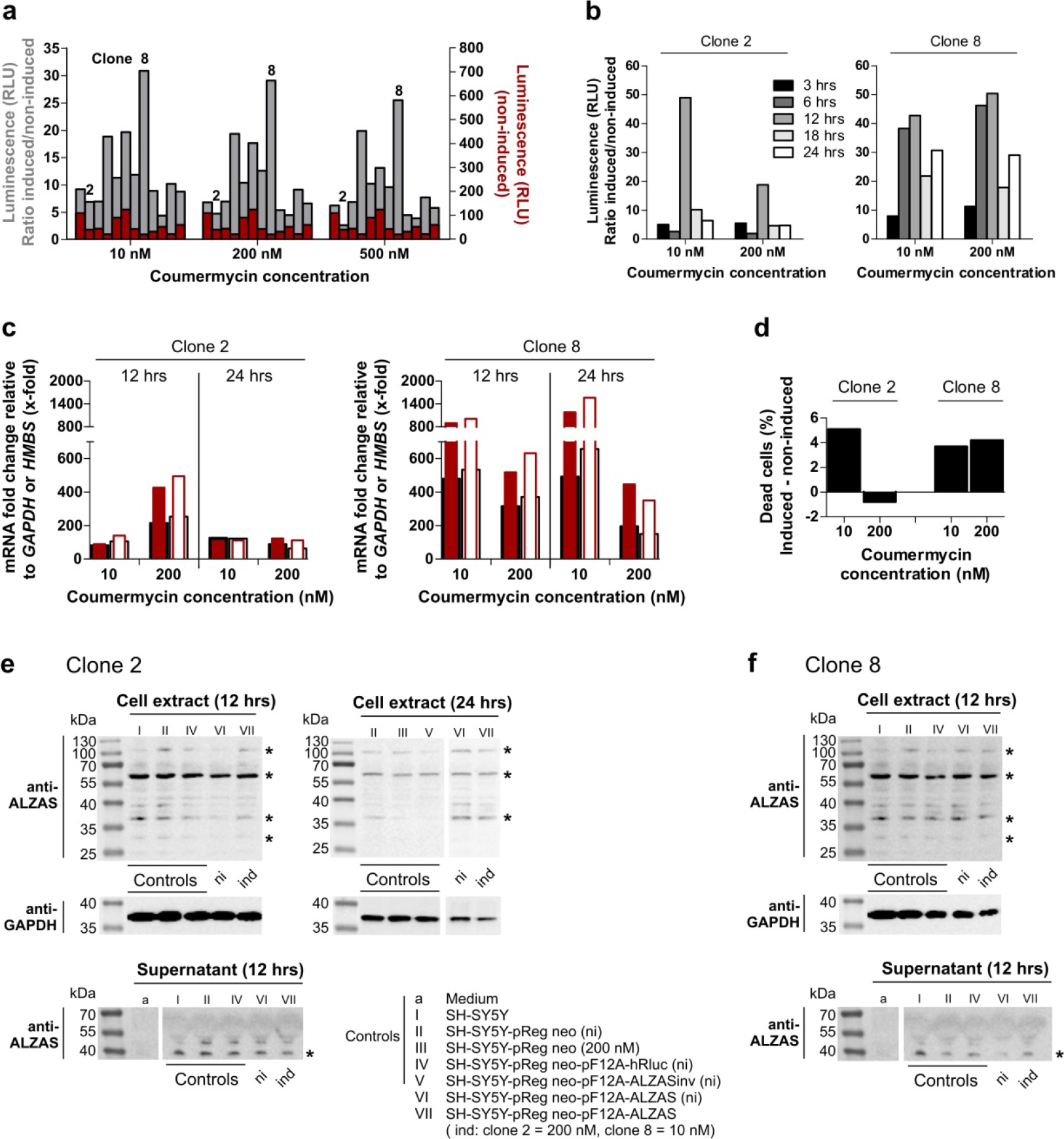
Gene expression analysis of *ALZAS* mRNA and protein level in human SH-SY5Y cells. (a) For a *Renilla* luciferase test expression, 12 different SH-SY5Y-pReg neo clones were transfected for 3 hrs with pF12A-hRluc. After transfection, coumermycin was added at a concentration of 10, 200 or 500 nM to induce luciferase expression. 24 hrs later, the activity of luciferase was measured. Red bars illustrate luciferase activity in the non-induced condition, gray bars illustrate the activity ratio between induced and non-induced samples (each condition normalized to no-template controls). Measurements were done once or repeated up to 3 times (value or mean is shown). (b) To analyze time dependence of *Renilla* luciferase expression, SH-SY5Y-pReg neo clone 2 and 8 were transfected for 3 hrs with pF12A-hRluc. Luciferase expression was induced with coumermycin. 3-24 hrs later, the luciferase activity was measured. Measurements were done once or repeated up to 3 times (value or mean is shown). (c, d) To analyze *ALZAS* (over)expression cells from SH-SY5Y-pReg neo, clone 2 or 8 were transfected for 3 hrs with pF12A-ALZAS before *ALZAS* expression was induced with coumermycin (clone 2: n = 2, clone 8: n = 1). (c) 12 or 24 hrs after induction, the *ALZAS* mRNA level was analyzed with 2-step RT-qPCR using normalization to *GAPDH* (filled bars) and *HMBS* (open bars). Relative changes in *ALZAS* expression compared to controls (mean of no-template and Rluc) are illustrated. A higher *ALZAS* mRNA level is presented by bars above 1. Black bars (filled/open) illustrate *ALZAS* expression measured with primer pair [a], red bars (filled/open) represent *ALZAS* mRNA measurement using primer pair [b]. Data is presented as mean determined from at least duplicate measurements. (d) 24 hrs after induction, a live/dead staining was performed. The number of dead cells compared to controls (described in material and methods) is illustrated. (e, f) SH-SY5Y-pReg neo clone 2 (e) or clone 8 (f) was transfected for 3 hrs with pF12A-ALZAS and followed by induction of *ALZAS* expression (n = 2). Controls (no-template, Rluc, *ALZAS*inv) were included. After 12 and 24 hrs, cell lysates were immunoblotted for ALZAS and GAPDH. For better illustration, vertical blot lanes were partially rearranged. Lanes originated from different blots are separated. Asterisks indicate protein bands interacting with anti-ALZAS antibody at about 30, 38, 60 and 100 kDa.
*ALZAS*inv: inverse *ALZAS* coding sequence, ni: non-induced, RLU: Relative Light Unit, Rluc: *Renilla* luciferase.

### Verification of the specificity of anti-ALZAS antibody for ALZAS protein and examine the binding of Aβ-antibodies to ALZAS protein

In the current study, we used a specifically produced anti-ALZAS antibody for all ALZAS protein detections. This antibody detects the same amino acid sequence (ct12) as the autoantibody found in human blood. To confirm the anti-ALZAS specificity, we performed IP followed by MS-based identification of ALZAS protein. IP-based purification of EGFP-ALZAS fusion protein generated in SH-SY5Y cells did not reach the required protein amount for MS measurement, although we employed cell lysates with the highest protein amount after 18 hrs (Supplementary Figure 3a). Hence, we applied *in vitro* translation of an EGFP-ALZAS fusion gene construct (T7-EGFP-ALZAS-polyA) using the TNT T7 Quick for PCR DNA System and TNT T7 Quick Coupled Transcription/Translation System (each Promega) as both kits provide an adequate amount of EGFP-ALZAS protein. The *in vitro* generated EGFP-ALZAS fusion protein (Figure 4a) was purified via GFP-binding beads prior to MS. In MS, altogether 117 fragmentation spectra were matched to 22 fully and semi-tryptic peptides from the EGFP-ALZAS construct; four of them were unique for ALZAS: LVFFAEDVGSNK, KQYTSIHHGVVEVGK, QYTSIHHGVVEVGK and LDCMFPSGN (Supplementary Figure S3b). In Western blot analysis, we detected the EGFP-ALZAS fusion protein with anti-GFP antibody as well as with the anti-ALZAS antibody and anti-beta Amyloid antibody recognizing amino acids 1-13 (Figure 4b). Further, we verified the identity of the EGFP-ALZAS fusion protein in a clinically applied, commercially available ELISA test for Aβ_42_ (INNOTEST β-AMYLOID_(1-42)_, Fujirebio; data not shown). However, tested polyclonal Aβ-antibodies interacting with Aβ_1-14_ or Aβ_22-35_ did not detect EGFP-ALZAS.

**Figure 4.**
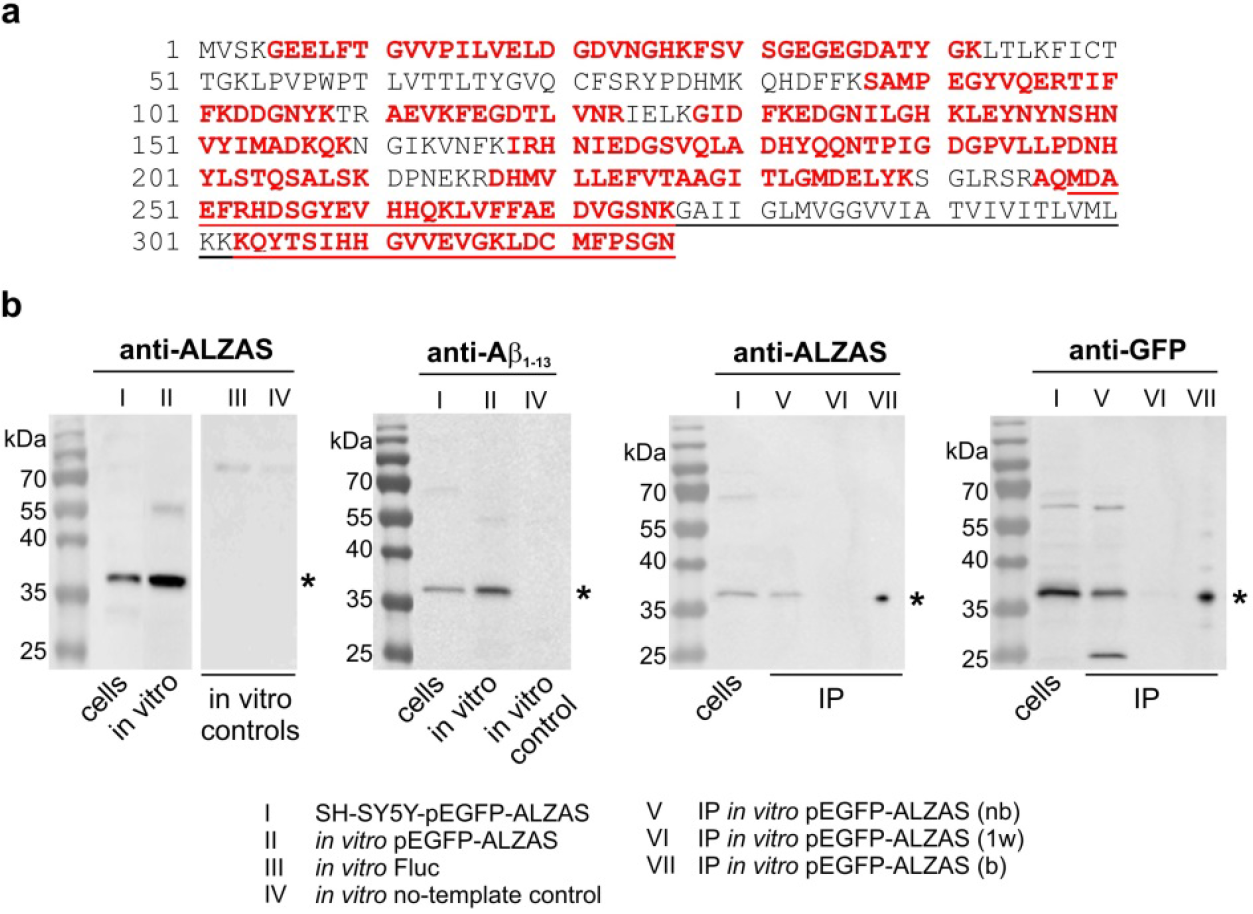
Analysis of anti-ALZAS specificity by MS and immunoblotting. (a) EGFP-ALZAS protein construct sequence. The amino acid sequence unique for ALZAS is underlined, MS-matched peptides are in red. (b) SH-SY5Y cells were transfected with pEGFP-ALZAS and 24 hrs later, the cell lysates immunoblotted for ALZAS, EGFP and Aβ_1-13_ (lane I). *In vitro* translation was performed for EGFP-ALZAS and controls (*Firefly* luciferase (Fluc), no-template) and immunoblotted for ALZAS and Aβ_1-13_ (lane II,III,IV). *In vitro* generated EGFP-ALZAS was applied for immunoprecipitation and aliquots of the non-bound fraction, first wash and bound fraction (each 10 μl) were immunoblotted for ALZAS and EGFP (lane V,VI,VII). For better illustration, vertical blot lanes were partially rearranged. Lanes originated from different blots are separated. Asterisks indicate EGFP-ALZAS fusion protein at about 36 kDa. nb: non-bound fraction, 1w: first wash fraction, b: bound fraction.

### Cell localization of the ALZAS protein

To identify the localization of the ALZAS protein in human, especially neuronal cells, we conducted immunocytochemical staining and analyzed them by fluorescence microscopy. We co-stained ALZAS (using anti-ALZAS antibody) with APP or markers for the cytoskeleton (F-actin), the endoplasmatic reticulum (protein disulfide isomerase, PDI), the Golgi apparatus (Golgi matrix protein of 130 kDa, GM130), mitochondria (Cytochrome C), lysosomes (Cathepsin) or endosomes (Ras-related in brain Rab5, Rab7). Fluorescence microscopy revealed a punctate pattern of ALZAS staining and a colocalization with the endoplasmatic reticulum (Figure 5a, b), APP (Figure 5a) or early endosomes (Rab5; Figure 5c), as indicated in yellow. An overlay with other cell organelles was not evident (Supplementary Figure S4). Further investigations in human tissue suggested a localization of ALZAS within the cell membrane as well as in cytoplasm based on Western blot analysis (data not shown).

No considerable differences were observed between SH-SY5Y control cells and *ALZAS* overexpressing SH-SY5Y-pReg neo-pF12A-ALZAS cells (clone 2 and 8). Cells of SH-SY5Y-pReg neo-pF12A-ALZAS clone 8 (non-induced) exhibited a remarkable staining with considerable accumulations around the nucleus and cell membrane (Figure 5b, indicated by arrows). Co-staining of endogenous ALZAS and the EGFP-ALZAS fusion protein in SH-SY5Y-pEGFP-ALZAS cells revealed no overlapping fluorescence signals for the two proteins (Figure 5d). Additionally, staining of ALZAS and organelles in SH-SY5Y-pEGFP-ALZAS cells exhibited no colocalization (Supplementary Figure S4b).

**Figure 5.**
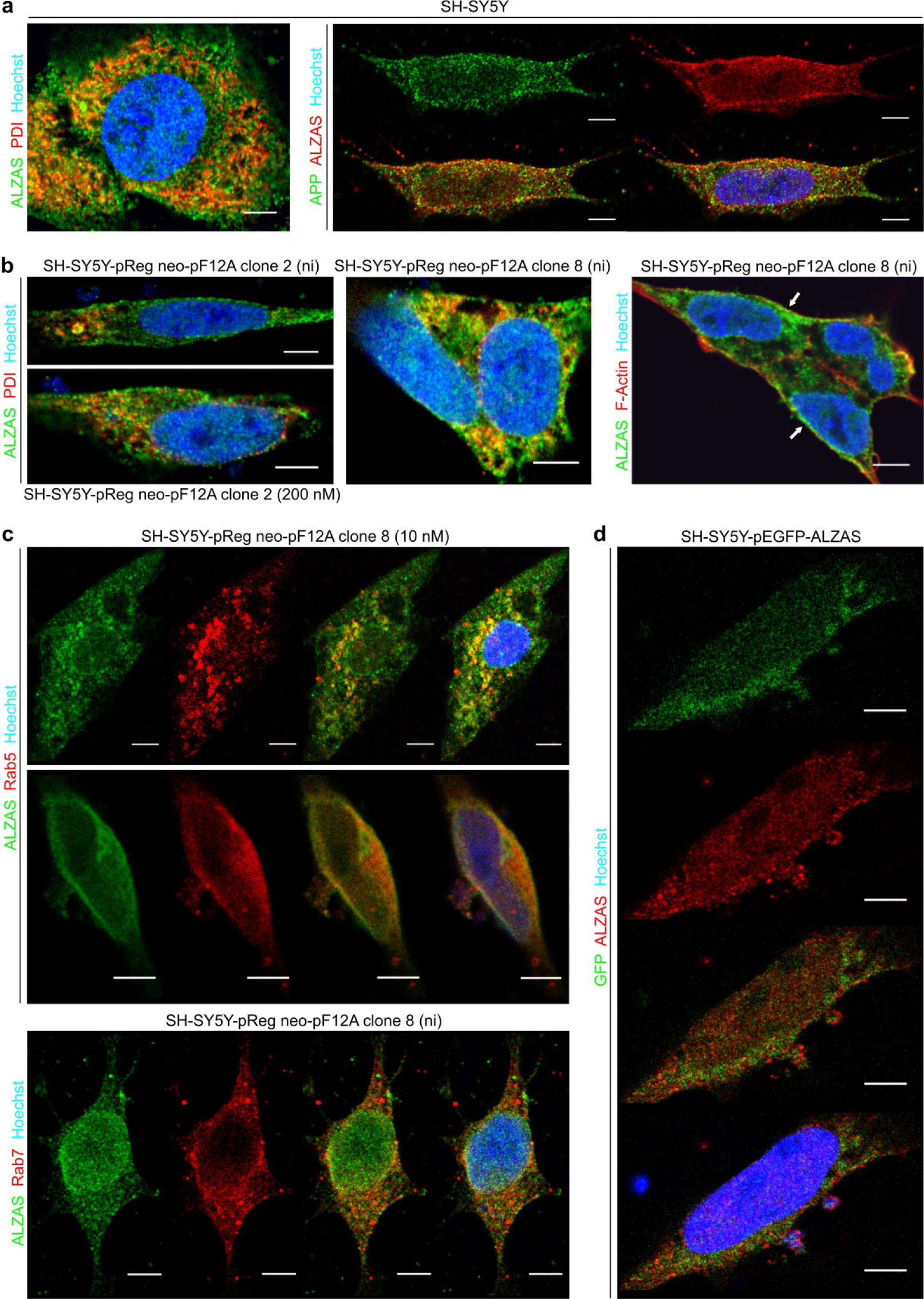
Cell localization of endogenous *ALZAS* protein, overexpressed ALZAS and EGFP-ALZAS fusion protein in human cells. (a) Neuroblastoma cells SH-SY5Y were immunocytochemically co-stained for ALZAS and PDI or APP and analyzed by confocal microscopy. Colocalization is indicated by yellow colour. (b, c) SH-SY5Y-pReg neo clone 2 and 8 were transfected for 3 hrs with pF12A-ALZAS and subsequently *ALZAS* expression was induced. 24 hrs later, the cells were immunocytochemically co-stained for ALZAS and (b) PDI or F-actin or (c) rab proteins (Rab5: early endosomes, Rab7: late endosomes) and colocalization (indicated by yellow) was determined by confocal microscopy. Arrows indicate potential accumulations of the ALZAS protein. (d) SH-SY5Y cells were transfected with pEGFP-ALZAS and 24 hrs later immunocytochemically labeled for ALZAS and EGFP and analyzed by confocal microscopy. Nuclear DNA was visualized with Hoechst 33342. Representative images are shown. Scale bar: 5 μm, ni: non-induced.

## Discussion

The *ALZAS* gene represents a special type of nested gene because it is located on the same DNA strand as the host gene, consists of exon and intron sequences of *APP* and occupies the same reading frame (Figure 1) (Bergmann and Preddie 1998). Mostly, eukaryotic nested genes, which are a specific class of overlapping genes, are encoded on the opposite DNA strand of the host gene (Gibson et al. 2005), either within a large intron (intronic nested genes) or a protein coding DNA sequence (non-intronic nested genes) (Kumar 2009). Just a third of nested genes are localized on the same DNA strand as the host gene (Yu et al. 2005). These different gene structures, especially the type and length of overlapping base pairs between nested and host gene, affect the relation of expression level of both genes (Ho et al. 2012). Rarely do host and nested gene exhibit the same expression level (Sanna et al. 2008). Accordingly, in the present study, we observed an approximately 1000-fold lower expression of *ALZAS* in tissues compared to *APP* (Figure 2d). *ALZAS* exhibited an ubiquitous transcription profile independent of sex and AD disease (Figure 2b, c) disconfirming initial hypotheses that *ALZAS* is either expressed solely in AD patients or overexpressed in AD brains (Bergmann et al. 1996; Kienzl et al. 2002; Kienzl et al. 2007). Furthermore, we confirmed a tissue-specific transcription of *ALZAS* as was already validated for *APP* (Kummer et al. 2002; Ge et al. 2004). Only 22% of protein coding human nested genes feature a tissue-specific expression (Yu et al. 2005; Ho et al. 2012). In blood of children, we verified the lowest *ALZAS* mRNA level. As different transcription profiles are ascribed to a stronger regulation of transcription (Viskochil et al. 1991; Yu et al. 2005; Ho et al. 2012), we assume that *ALZAS* is subject to a dual control system under normal conditions since it is part of two precursor mRNAs (its own and from *APP*). A separate promoter region for *ALZAS* is postulated to be located within intron 15 of the *APP* gene, consisting of heat shock and estrogen responsive elements (Bergmann and Preddie 1998). Other regulatory post-transcriptional mechanisms exist includes RNA accumulation prior to degradation (Henikoff and Eghtedarzadeh 1987) or the stabilization of mRNA through binding of natural antisense transcripts (NATs) (Werner and Sayer 2009) which might apply for *ALZAS*. However, further investigations are required to identify specific *ALZAS* transcription factors. Simultaneous transcription of *ALZAS* and *APP* may trigger interferences like steric hindrance of both polymerases and/or *ALZAS* regulator proteins (Gibson et al. 2005).

Unfortunately, in the present study, a linked expression of *ALZAS* and *APP* could neither be confirmed nor disproved.

We obtained exceptional data in human placenta samples where we measured an *ALZAS* mRNA concentration similar to that in brain and other peripheral tissues (data not shown). Usually, in humans, the placenta-transcriptome is unique (Kim et al. 2012). To underscore our tissue-specific expression of *ALZAS* and to investigate a statistical significance for *ALZAS* expression pattern, we suggest that other tissues and/or a much bigger amount of different tissues should be examined. Further, additional studies are required to verify a potential age- and disease-dependent transcription of *ALZAS*.

In the current study, analyses of *ALZAS* translation and protein amount in various tissues were performed for the first time, except for an enzyme-linked immunosorbent assay (ELISA) from human serum, which revealed an elevated amount of ALZAS protein in late clinical AD stages (Kienzl et al. 2007). Our investigations in the brain (see Table S1) revealed no considerable differences in protein signals of non-AD controls and AD patients (Figure 2e). However, we did observe different sizes and intensities of protein bands interacting with anti-ALZAS antibody for each tissue. We postulate that the ALZAS protein (calculated size of monomer is about 9 kDa) develops stable oligomeric structures which are part of ADDLs, as was established for Aβ-peptides (Klein et al. 2001). Aβ-ADDLs are heterodisperse molecules consisting of soluble oligomers of different sizes (dimers to up to 24mers) whose amount is elevated in AD-affected brain regions (Klein 2002). While Aβ-ADDLs are mainly constituted of trimers to up to pentamers (Chromy et al. 2003), our Western Blot analyses suggest that ALZAS might form hexamers in brain and probably hexamers and trimers in liver. The existence of stable oligomer formation, even under reducing conditions of denaturing SDS-PAGE, has been determined for Ae-peptides (Enya et al. 1999) and is also probable for ALZAS which contains the entire Aβ_42_-sequence. It might also be possible that ALZAS form oligomeric structures along with Aβ-peptides. In previous immunohistochemical studies, ALZAS could be detected in amyloid plaques using anti-ALZAS antibody (Kienzl et al. 2002). The aggregation of Aβ_42_-peptides is ascribed either to (I) the hydrophobic C-terminus (amino acids 17 to 42) (Jarrett et al. 1993), (II) β-sheet conformation (Liao et al. 2007), (III) pH value, (IV) protein concentration (Lansbury 1999) or (V) stability of the native, soluble protein (Dobson 2001). The shape of At-oligomers is further influenced by their soluble or membrane-bound nature. Soluble Aβ-species appear in monomers to hexamers, while membrane-associated Ae-peptides form larger aggregates (Johnson et al. 2011). To confirm our considerations of ALZAS aggregation we recommend the analysis of the different sized ALZAS oligomers. Moreover, it is necessary to examine a bigger variety of organs for their ALZAS protein pattern.

A comparison of transcription and translation levels of *ALZAS* in human tissues revealed no correlation between mRNA and protein levels, which are described for many other genes, as well. Weakly expressed genes mostly possess no correlation between mRNA and protein level (Maier et al. 2009). Contrary to the *ALZAS* mRNA level, we detected prominent protein signals in liver (Figure 2f) and none in skeletal muscle of an AD patient (data not shown). Previously, attenuated levels of Aβ-peptides have been observed in liver of AD patients (Roher et al. 2009). Here we postulate that, not only APP but also ALZAS represents a source of Aβ-peptides and the probably high amount of ALZAS protein in AD liver may evoke an increased Aβ_42_-concentration, being rediscovered as amyloid plaques in AD brain. Since non-demented controls also exhibit an elevated Ae-concentration in liver, it is unclear if the liver sequesters or is the source of Aβ-peptides (Sagare et al. 2011).

Investigations concerning the discrepancy between mRNA and protein amount of *ALZAS* are necessary to understand the regulation of *ALZAS* gene in detail. It might be influenced by *APP* mutations in exon 16/17 (Bergmann and Preddie 1998) prior to post-transcriptional regulation, for instance, RNA interference (Meister 2007) or NATs, which influence mRNA processing, RNA transport and tissue-associated translation (Werner and Sayer 2009). Moreover, *ALZAS* gene regulation could be affected at post- or co-translational level because the obviously impaired ubiquitin proteasome system (UPS) in AD patients (Oddo 2008) might fail in the clearance of ALZAS protein. While several mechanism causing tissue- or temporal-specific translation were identified for the host gene *APP* (Kao et al. 2004; Delay et al. 2012; Fan et al. 2013), our investigation of induced *ALZAS* overexpression in the human neuroblastoma cell line SH-SY5Y did not lead to considerable alterations in ALZAS protein levels (Figure 3e, f, Supplementary Figure 2c). A greatly enhanced *ALZAS* transcription, from >200- up to >500-fold depending on the SH-SY5Y-pReg neo clone, was, however, verifiable (Figure 3c). Nevertheless, *ALZAS* mRNA expression was still far below *APP*.

In contrast to tissue samples, we detected different protein bands interacting with anti-ALZAS antibody in the SH-SY5Y cells (Figure 3e, f). We assume that this pattern reveals different oligomer formations of ALZAS. This observation might correlate with the size of Ah-oligomers which vary in cell specific manner (Klein 2002). The apparently unchanged protein amount in induced *ALZAS* overexpressing cells compared to controls (Figure 3e, f, Supplementary Figure S2c) is in accordance with the facts that no considerable increase in cell death (Figure 3d) and no higher intensity of ALZAS fluorescence signals (Figure 5b, c) were observed upon overexpression. Since anti-ALZAS antibody could not distinct between endogenous and overexpressed (induced condition) ALZAS protein, we have estimated to find stronger signals in *ALZAS* overexpressing cells through higher ALZAS protein amount. Furthermore, we did not detect ALZAS protein in detached *ALZAS* overexpressing SH-SY5Y cells (data not shown).

However, since the *Renilla* luciferase assays unraveled an inductor concentration-dependent translation (Figure 3a, b), it might be assumed that an enhanced amount of ALZAS protein could have negative effects on human cell metabolism. Thus, post-transcriptional and/or post-translational regulation mechanisms limit the amount of ALZAS protein in overexpressing cells. This apparently strong, regulation of *ALZAS* implies a considerable function of the protein in cell metabolism, as expected for all human nested genes (Yu et al. 2005). To date, it is a general belief that ALZAS is a transmembrane protein which could be integrated in the membrane similar to or instead of APP because ALZAS possesses the whole transmembrane domain of APP (Figure 1) (Jellinger et al. 2008). As we were able to detect ALZAS in the supernatant of overexpressing cells (Figure 3e, f), we speculate that it might have crossed the membrane by secretion. For AA-peptides, it has been validated that they are secreted followed by the extracellular deposition (Dobson 2001). However, in AD, Aβ-release may occur initially after cell death (Recuero et al. 2004).

Amphiphilic properties are affirmed for ALZAS and also for peripheral membrane proteins which are not stably embedded in the membrane and thus can be liberated (Bhardwaj et al. 2006). Our investigation, using a protocol for fractionated protein isolation, indicated that ALZAS is localized in the membrane and the water-soluble protein fraction (data not shown). Further hints that ALZAS might be a peripheral membrane protein, for example, in vesicles, were found in immunocytochemical staining. The abundant ALZAS staining near the cell membrane and around the nucleus (Figure 5a-c) showed a predominant overlay with Rab5 protein (Figure 5c), a peripheral membrane protein in early endosomes involved in cellular signal transduction (Chavrier et al. 1990). An overlay of ALZAS and APP was observed more in the cell membrane/cell surface (Figure 5a), where APP is spliced in the non-amyloidogenic pathway (s-cleavage) (Kojro and Fahrenholz 2005). On the contrary, the amyloidogenic cleavage occurs mainly in endosomes (Vassar et al. 1999). However, the EGFP-ALZAS fusion protein showed a divergent localization to endogenous ALZAS (Figure 5d) which might indicate that, upon fusion of EGFP to the N-terminal, a crucial transport signal of ALZAS is lost. In contrast, the EGFP-Rab5 fusion protein showed colocalization with endogenous Rab5 in cytoplasmic vesicles (Hoffenberg et al. 2000). However, similar to ALZAS, Rab5 is ubiquitously expressed and undergoes aggregation. An overexpression of Rab5 achieved no significant enhanced membrane staining (Chavrier et al. 1990) which is analogous to our immunocytochemical staining in *ALZAS* overexpressing SH-SY5Y cells.

In addition to Rab5, the APP protein and the I-amyloid cleaving enzyme 1 (BACE1), mediating β-secretase cleavage of APP, are prevalently localized in early endosomes (Kinoshita et al. 2003), and more precisely, in the membrane structure termed lipid raft, which represent the primary place of cellular signal transduction (Fantini et al. 2002). Several proteins encoded by nested genes are involved in cell-cell communication and our studies suggest that ALZAS may play a similar role. Comparing the cellular functions of nested and host genes, a correspondence is rarely observed (Yu et al. 2005). However, APP may perform a regulatory function as a receptor or in cell adhesion (Sondag and Combs 2004; Sosa et al. 2013). Interestingly, prevention of APP (-cleavage did not result in endosome dysfunction which is typical for patients in early AD stages (Jiang et al. 2010).

In addition to extensive transcription, translation and cell localization studies, we could, for the first time to our knowledge, prove the ALZAS amino acid sequence and the specificity of anti-ALZAS antibody produced using the antigenic peptide ct12 that act as immunogenic sequence in humans. Using EGFP-ALZAS fusion protein produced *in vitro*, we confirmed the specific attachment of anti-ALZAS antibody to ALZAS, while the tested monoclonal Aβ_1-13_-antibody recognized both ALZAS (Figure 4b) and Aβ_40_. Moreover, the tested monoclonal Aβ_37-42_-antibody, a commercially available ELISA test for diagnosis of AD, recognized both the Aβ_42_ and the EGFP-ALZAS fusion protein, revealing that this assay is not specific for Aβ_42_ but may as well detect ALZAS. However, we could not detect EGFP-ALZAS with tested polyclonal Aβ-antibodies interacting with Aβ_1-14_ or Aβ_22-35_, indicating a conformational divergence of epitopes. Taken together, immunodetection of β,-peptides in former studies come under suspicion of co-detecting ALZAS.

Considering that highest ALZAS autoantibody concentrations were observed in patients with early AD, we agree with earlier assumptions that ALZAS could serve as an indicator for AD development and may be a diagnostic blood marker (Bergmann and Preddie 1998). With respect to AD-specific expression of *ALZAS*, further investigation are necessary to confirm the presence of ALZAS in amyloid plaques (Kienzl et al. 2002) and its function in the human. In conclusion, the present study demonstrates that ALZAS has the potential to be a novel and crucial factor in AD pathology.

## Acknowledgements

We thank Prof. Jellinger (Institute of Clinical Neurobiology, Vienna) and E. Kienzl (Technische Universität Wien); Prof. V. Holthoff-Detto, Dr. Plötze, Ms Rings (Technische Universität Dresden) for tissue samples; Dr. J. Bergmann (Universitätsklinikum Hamburg-Eppendorf) and Dr. S. Kretschmer (Technische Universität Dresden) for advice and assistance in IP, Prof. van Pée (Technische Universität Dresden) and Dr. Francisco Pan-Montojo (Ludwig-Maximilians-Universität München) for helpful discussions and Dr. W. Mages (University of Regensburg) for careful reading of the manuscript. We would like to thank the Light Microscopy Facility (especially S. Tulok, Dr. A. Walther) for confocal imaging, the Mass Spectrometry Facility of the MPI-CBG, Dresden (especially Dr. A. Shevchenko) and Prof. S. Bergmann and D. Mielsch (Technische Universität Dresden) for Aβ_42_-ELISA sample measurement. This work was partially supported by the Association of Friends and Sponsors of TU Dresden e.V. and the Affirmative Action Plan for Women at Technische Universität Dresden. The authors indicate no potential conflicts of interest.

